# Gestational pertussis vaccination and the infant’s cellular immune response against whole-cell pertussis vaccine in the first year of life

**DOI:** 10.1101/697722

**Authors:** Carolina Argondizo-Correia, Lourdes Rehder de Andrade Vaz-de-Lima, Elaine Uchima Uehara, Eder Gatti Fernandes, Helena Keico Sato, Ana Paula Sayuri Sato, Expedito José de Albuquerque Luna, Euclides Ayres de Castilho, Cyro Alves de Brito

## Abstract

Pertussis resurgence worldwide calls for new prevention strategies, as the recently incorporated vaccine booster dose during pregnancy, whose aim is to protect newborns from infection. In Brazil, maternal Tdap vaccination is recommended since 2014, and we reported that this strategy promotes high transplacental transfer of anti-PT IgG and it is effective in protecting infants early in life. Young children are the most susceptible group and with higher mortality rates, however, it is not well known whether the elicited anti-pertussis maternal antibodies could influence in the children’s immune responses further in life, especially after their own vaccination against pertussis. Considering this scenario, we conducted a study with children born to mothers who either received or not the booster dose during pregnancy, after their primary pertussis vaccination, in order to investigate the first impact of maternal immunisation on the response to infant immunisation regarding the cellular immune response, while comparing with data from the literature. As transfer of maternal antibodies could result in attenuation of the immune response to vaccination in infants, this study performed to determine whether higher levels of maternal antibodies could influence in the immune response of infants to the whole-pertussis vaccination series. Results showed no difference in cytokine production between groups, a first suggestion that maternal vaccination may not interfere with recognition and cellular response generation to vaccination. This data, together with humoral immunity and epidemiological studies, is important for the implementation of maternal immunisation strategies nationwide and will contribute to assure public policies regarding vaccination schemes.

**Importance:** Pertussis, or whopping cough, is a respiratory infectious disease caused by a bacterial agent, resulting in violent coughs and possibly death in vulnerable groups, such as young children and neonates. It is known that pregnant mothers transfer antibodies to their developing foetuses for protection in early life, however anti-pertussis antibodies are not highly detected in young children. Thus, a pertussis maternal vaccination was implemented to increase maternal anti-pertussis antibodies levels in pregnant women and therefore the transference to the foetus. However, maternal antibodies can also interfere in the child immune response in the first months of life. The significance of our research is in analysing the cellular immune response of children born from pertussis-vaccinated mothers, which will give a first glimpse on how maternal antibodies could modulate the child’s response to pertussis in early life.

## Introduction

Pertussis is a contagious respiratory disease caused by the *Bordetella pertussis* bacteria [1]. Despite high vaccination coverage, it remains an important public health problem, re-emerging in several countries every several years [2,3]. It has a high rate of morbimortality in young children, and estimates from the World Health Organization suggest that ~50 million pertussis cases and 300,000 deaths occurred annually, mostly in children under five years of age [4]. In Brazil, between 2010 and 2014, over 22,000 people were infected [5], most cases in children younger than 1 year of age [5,6].

The current childhood vaccination schedule includes three doses of the diphtheria, tetanus and whole cell pertussis vaccine, combined with *Haemophilus influenza* b and hepatitis B (DTwP-Hib-HBV) in the first year of life, at 8, 16 and 24 weeks of age, leaving neonates without specific protection. So, a dose of tetanus, reduced diphtheria and acellular pertussis vaccine (Tdap) during pregnancy was proposed to address this immunity gap in young infants and promote a higher transplacental transfer of maternal antipertussis antibodies (MatAb) to the foetus, resulting in improved protection during the neonatal period and until their own vaccination scheme is completed [7].

Since 2012, this has been recommended in the United States, United Kingdom and Australia [8], and since November 2014 in Brazil [9]. Recently, we have shown Tdap maternal vaccination promotes high titers of anti-PT IgG in newborn [10], also this strategy showed a vaccine effectiveness of 82.6% in protecting infants aged <8 weeks from pertussis in Brazil [11]. However, until now, little is known whether it may affect the subsequent childhood vaccination, which uses whole-cell pertussis (wP) [12], while other countries recommend the acellular pertussis (aP) version [13]. There is also a concern that in addition to promoting protection to young infants, high MatAb titres could interfere or attenuate the immune response to the primary childhood pertussis vaccination series [14–16].

The humoral response is not the only responsible for the protection against infection, as well as there are no correlates of protection for serological levels of antipertussis antibodies [17]. Several studies show that the cellular immune response is required for effective clearance of infection from the respiratory tract and disease prevention, through effector mechanisms mediated by IFN-γ and IL-2, which are mostly produced by the T helper lymphocyte 1 (Th1). This cell type is described as having a more inflammatory profile, opposed to Th2 cells, characterized by the production of IL-4, IL-5 and IL-13. Some authors also relate the presence of IL-17 in the mechanism against *B.pertussis in vivo*, a cytokine produced by Th17 cells [18–23].

In light of this information, we sought to analyse the production of effector cytokines that could imply the activation of the infants’ cellular immune response against *B. pertussis* antigen *in vitro*, in the context of the presence of maternal anti-pertussis antibodies.

## Material and Methods

### Study Design and Participants

This study included 43 children around 7 months of age, born either to mothers vaccinated during pregnancy with a Tdap boost vaccine during the third trimester of pregnancy (n=33) or mothers who did not get vaccinated (n=10). This cohort is derived from a larger cohort described elsewhere [10]. Mothers were followed up after parturition until their children received the primary pertussis vaccination series, composed by three DTwP-Hib-HBV doses (produced by Bio-Manguinhos/Fiocruz), at approximately 2, 4 and 6 months of age. The protocol was approved by the Ethics Committee from the Adolfo Lutz Institute (CAAE: 37581114.0.0000.0059).

### Sample Processing and In Vitro Stimulation

5 mL of heparinized venous peripheral blood samples were collected by nurses from young children around one month after the third DTwP dose.

Samples were diluted in culture medium (RPMI 1640 [Gibco, Massachusetts/USA] supplemented with 2 g of NaHCO_3_, 10 mL of nonessential amino acids [Gibco], 40 mg of gentamycin and 10 mL of 200 mM L-glutamine [Sigma, Missouri/USA]). Peripheral blood mononuclear cells (PBMC) were isolated using a Ficoll-Paque Plus 1440 gradient (GE Healthcare, Uppsala/Sweden), according to the manufacturer’s instructions. Cells were washed and resuspended in culture medium, counted and rested overnight (~16 h, 1×10^6^ cells/mL) in 48-well flat-bottom plates at 37°C in a 5% CO_2_ incubator. On the following day, cells were stimulated with 2 μg/mL phytohemagglutinin (PHA) for 48 h [Sigma, Missouri/USA] or 5 μg/mL of inactivated pertussis toxoid (PT) for 120 h. After culture, cells and supernatant were collected and stored at −80°C for later assays. Cell pellets were stabilized using RNAlater (Sigma) and phosphate buffered saline (21.02 g of Na_2_HPO_4_, 7.16 g of NaH_2_PO_4_.H_2_O, 164.64 g of NaCl and reverse-osmosis H_2_O for 1 L) until total RNA extraction.

### Cytokine Detection

Frozen supernatants were thawed at room temperature, and cytokines were measured using both *Cytometric Bead Array* Th1/Th2/Th17 kit (BD Biosciences, New Jersey/USA) and LEGENDplex *Human Th Cytokine Panel* (13-*plex*) kit (BioLegend, California/USA), according to the manufacturers’ instructions, in order to compare results and analyse a larger range of cytokines. Both curve and samples were read in a BD LSRFortessa flow cytometer (BD Biosciences). Minimum detection limits were 4.88 pg/mL (BD) and 2.4 pg/mL (BioLegend). Corresponding analyses were performed with FCAP Array software v3.0 (BD Biosciences) and LEGENDplex Data Analysis software (BioLegend).

### RNA Extraction and Quantitative Polymerase Chain Reaction (qPCR)

RNA extraction was performed using Qiagen’s RNeasy Mini kit (Qiagen, Hilden/Germany), following the manufacturer’s instructions. To remove any traces of DNA, a DNA digestion step was performed using an RNAqueous Mini Kit (Ambion, Massachusetts/USA). RNA purity and quantity were evaluated by spectrophotometry in a Nanodrop (Thermo Scientific, Massachusetts/USA). RNA was reverse transcribed to cDNA using an iScript cDNA Synthesis Kit (Bio-Rad, California/USA) according to the manufacturer’s instructions.

In an attempt to identify different cell types, the chosen genes were the transcription factor *RORC2* and the cytokine gene *IL17A*, representing Th17 cells, the transcription factor *GATA3* and the cytokine gene *IL4* representing Th2 cells, and the cytokines *IFNG* for Th1 and *IL10* for regulatory response [24]. qPCR assays were run for each sample containing cDNA as a template, specific forward (5’-3’) and reverse (5’-3’) primers, respectively:

*RORC2:* TGGAAGTGGTGCTGGTTAGGA/AAGGCTCGGAACAGCTCCAT; *GATA3*: AGATGGCACGGGACACTACCT/TAATTCGGGTTCGCTTCCG; *IFNG*: GTTTTGGGTTCTCTTGGCTGTTA/AAAAGAGTTCCATTATGCGCTACATC; *IL17A*: GACTCCTGGGAAGACCTCATTGG/CTTGTCCTCAGAATTTGGGCATCC; *IL4*: ACAGCCTCACAGAGCAGAAGACT/TGTTCTTGGAGGCAGCAAAGA; *IL10*: CAGGGCACCCAGTCTGAGAA/CACATGCGCCTTGATGTCTG; *GAPDH*: GAAGGTGAAGGTCGGAGT/GAAGATGGTGATGGGATTTCCA; and SYBR Green PCR Master Mix (Life Technologies, Massachusetts/USA) on an Applied Biosystems 7500 Real-Time PCR System (Applied Biosystems, California/USA). *GAPDH* gene expression was used as a control to normalize the data. Results were represented as fold change given by the equation described by Livak and Schmittgen [25].

### Statistical Analysis

Descriptive and inferential statistical analyses were performed with GraphPad Prism software (version 5.0). Nonparametric Mann-Whitney, Kruskal-Wallis, Friedman, Dunn and Wilcoxon tests were used for inter- and intragroup analyses, and Spearman’s correlation coefficient was used for demographic and experimental data. We used α=5%, 95% confidence interval, level of significance=5% and 2-sided tests.

## Results

The study population consisted of young children born from women who received (VC) or not (NVC) a Tdap boost dose during pregnancy (Table 1). Samples were randomly chosen from a larger cohort described by Vaz-de-Lima et al. [10]. The groups presented a similar and representative demographic profile when compared to the larger cohort.

**Table 1.**
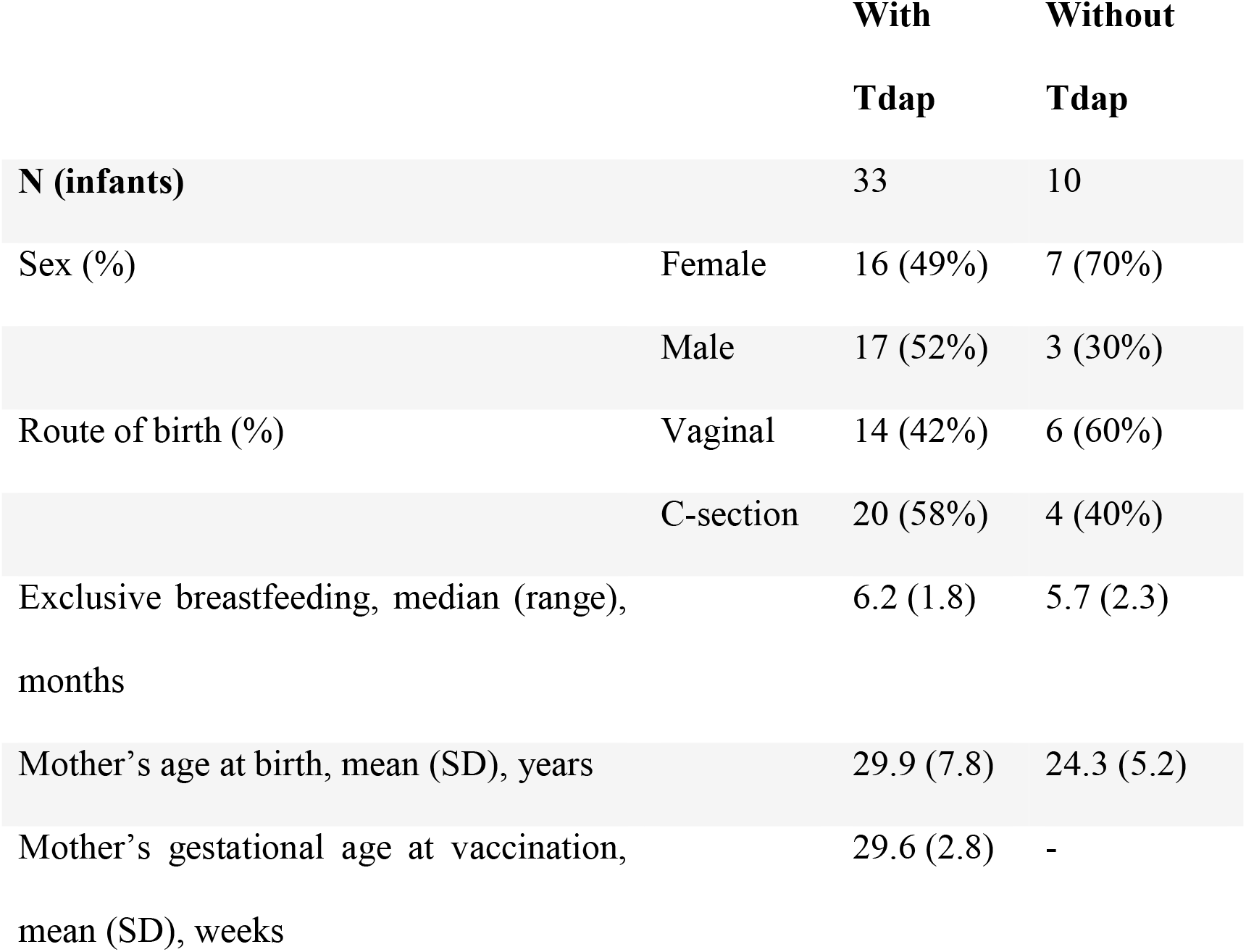
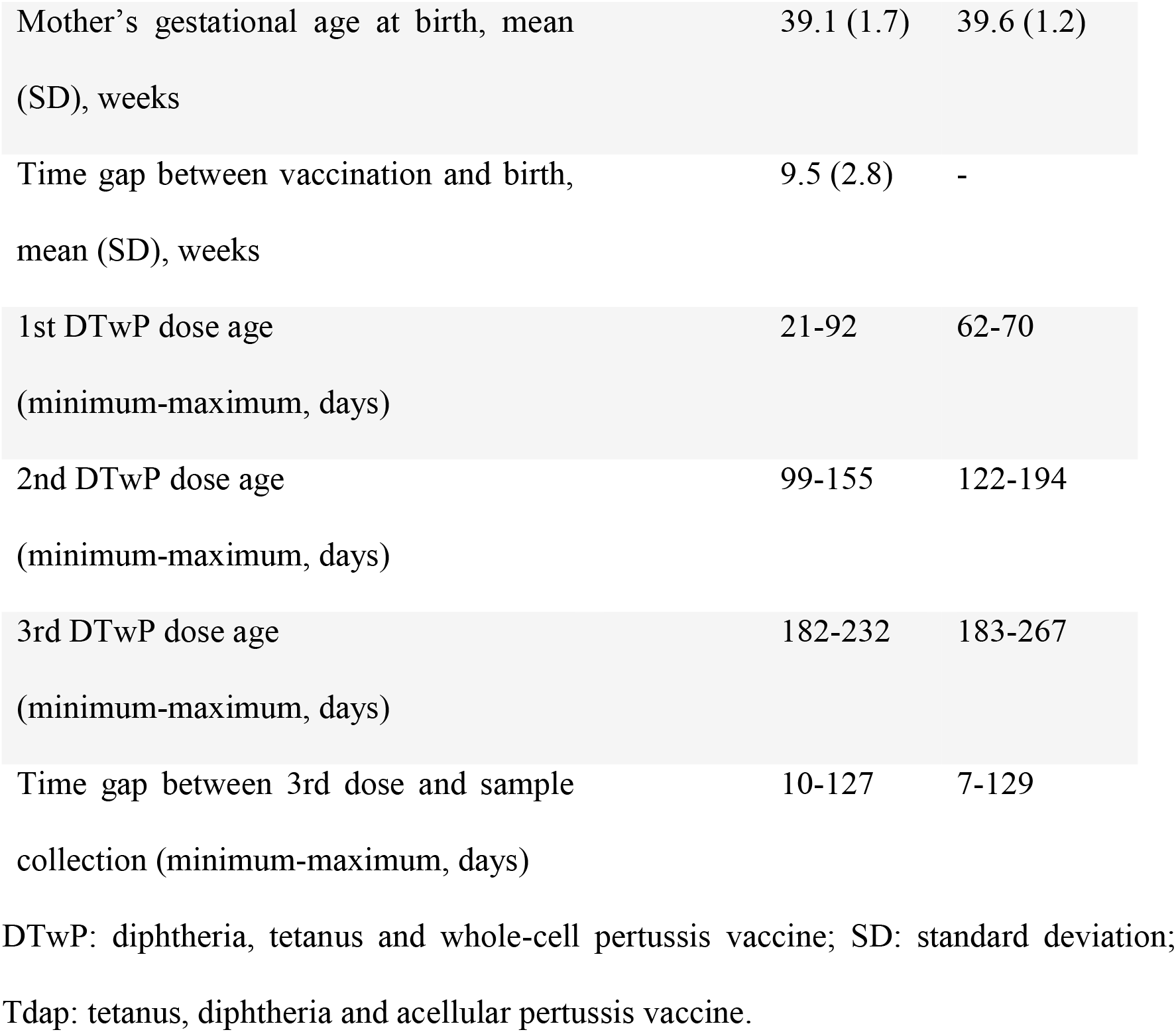
Information about the study groups

|  |  | **With**<br>**Tdap** | **Without**<br>**Tdap** |
| --- | --- | --- | --- |
| **N (infants)** |  | 33 | 10 |
| Sex (%) | Female | 16 (49%) | 7 (70%) |
|  | Male | 17 (52%) | 3 (30%) |
| Route of birth (%) | Vaginal | 14 (42%) | 6 (60%) |
|  | C-section | 20 (58%) | 4 (40%) |
| Exclusive breastfeeding, median (range),<br>months |  | 6.2 (1.8) | 5.7 (2.3) |
| Mother's age at birth, mean (SD), years |  | 29.9 (7.8) | 24.3 (5.2) |
| Mother's gestational age at vaccination,<br>mean (SD), weeks |  | 29.6 (2.8) | - |

|  |  |  |
| --- | --- | --- |
| Mother's gestational age at birth, mean (SD), weeks | 39.1 (1.7) | 39.6 (1.2) |
| Time gap between vaccination and birth, mean (SD), weeks | 9.5 (2.8) | - |
| 1st DTwP dose age (minimum-maximum, days) | 21-92 | 62-70 |
| 2nd DTwP dose age (minimum-maximum, days) | 99-155 | 122-194 |
| 3rd DTwP dose age (minimum-maximum, days) | 182-232 | 183-267 |
| Time gap between 3rd dose and sample collection (minimum-maximum, days) | 10-127 | 7-129 |

DTwP: diphtheria, tetanus and whole-cell pertussis vaccine; SD: standard deviation;
Tdap: tetanus, diphtheria and acellular pertussis vaccine.

In order to evaluate whether Tdap vaccination during pregnancy could induce modification in the cellular immune response of children, we evaluated a few related genes and cytokines that could represent the main cells in the cellular immune response, herein described as Th1, Th2 and Th17.

Transcriptional profile (Figure 1) showed no difference between groups, though the median levels of *GATA3*, *IL10* and *IFNG* were higher in children born to nonvaccinated mothers. When comparing the normalized expression of both basal and stimulated condition between groups, NVCs had lower levels for the basal condition concerning *GATA3* (p=0.0279) and *IFNG* (p=0.0290) expression. In the stimulated condition, the differences disappear, so the higher median level in the fold change analysis can indicate that in order to achieve the same expression upon PT stimulation, NVCs must have a greater increase in the expression of *GATA3* and *IFNG* due to the lower basal expression. To verify whether the results found in gene expression reflected the cytokine production, cytokines from Th1, Th2 and Th17 profiles were quantified via flow cytometry (Table 2). PHA stimulation was used as a positive control in a few samples for the main cytokines for the 3 Th profiles, due to samples and kit availability.

**Table 2.**
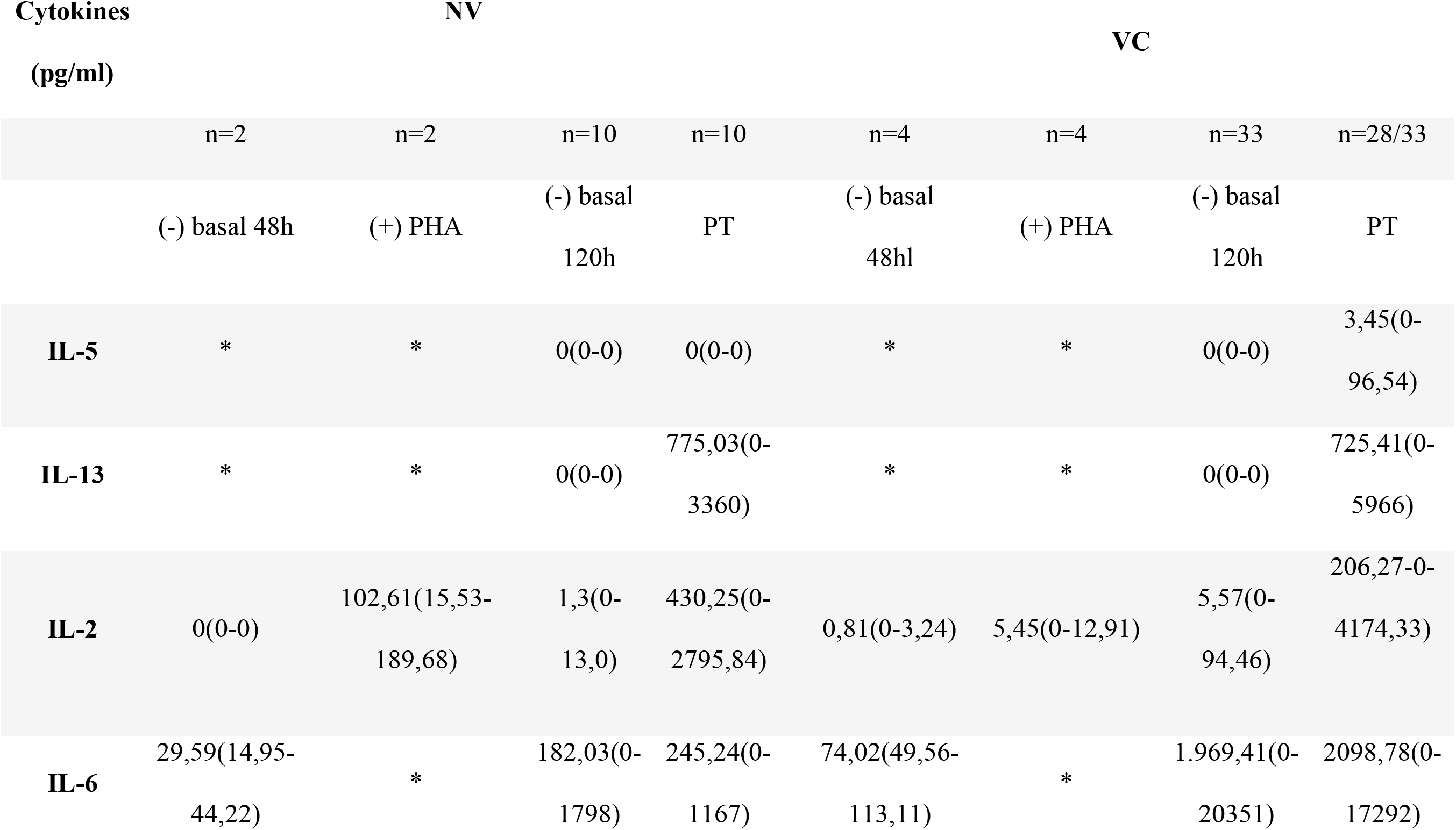
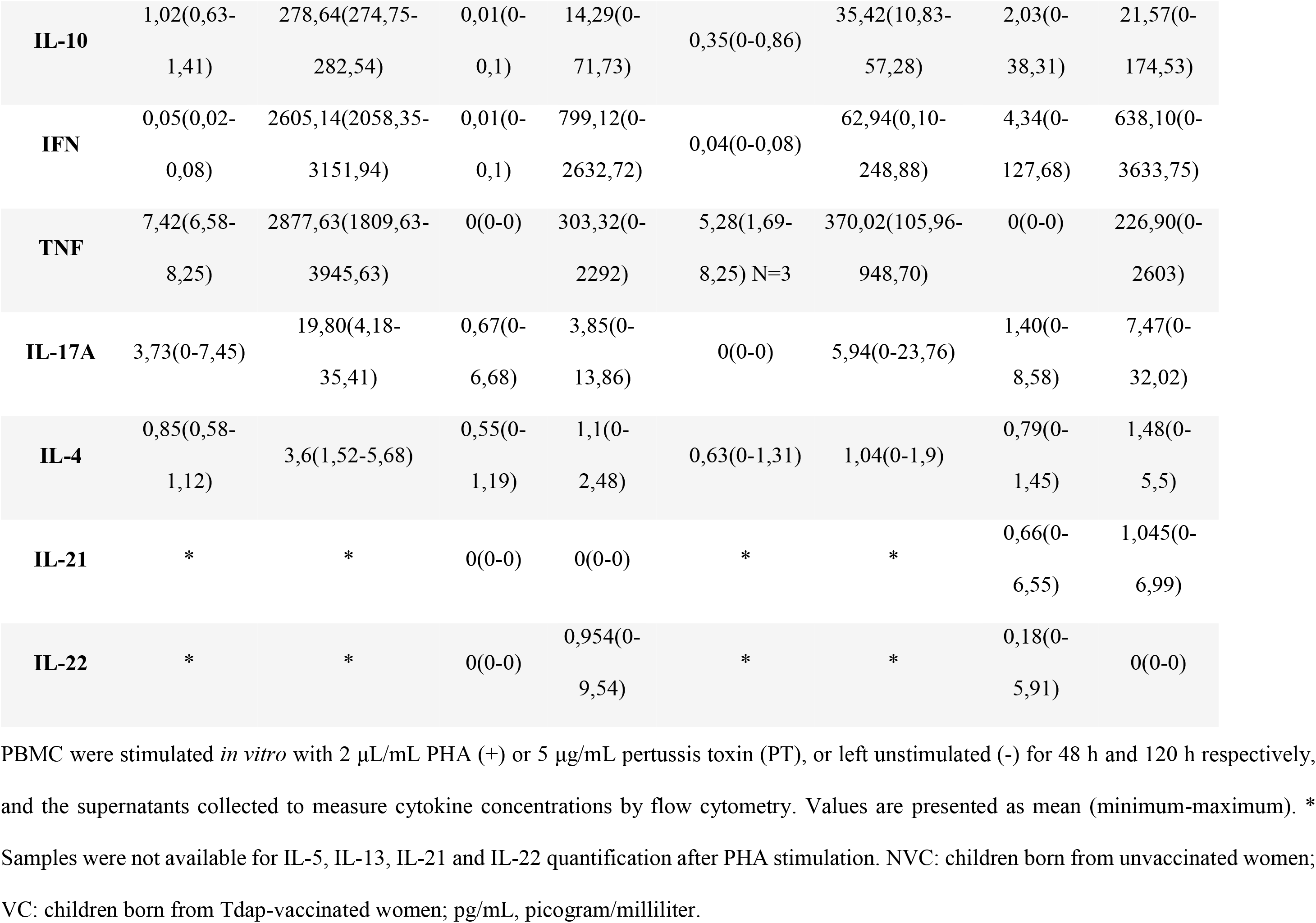
Cytokine concentration in culture supernatant of peripheral blood mononuclear cells (PBMC) from infants.

**Figure 1:**
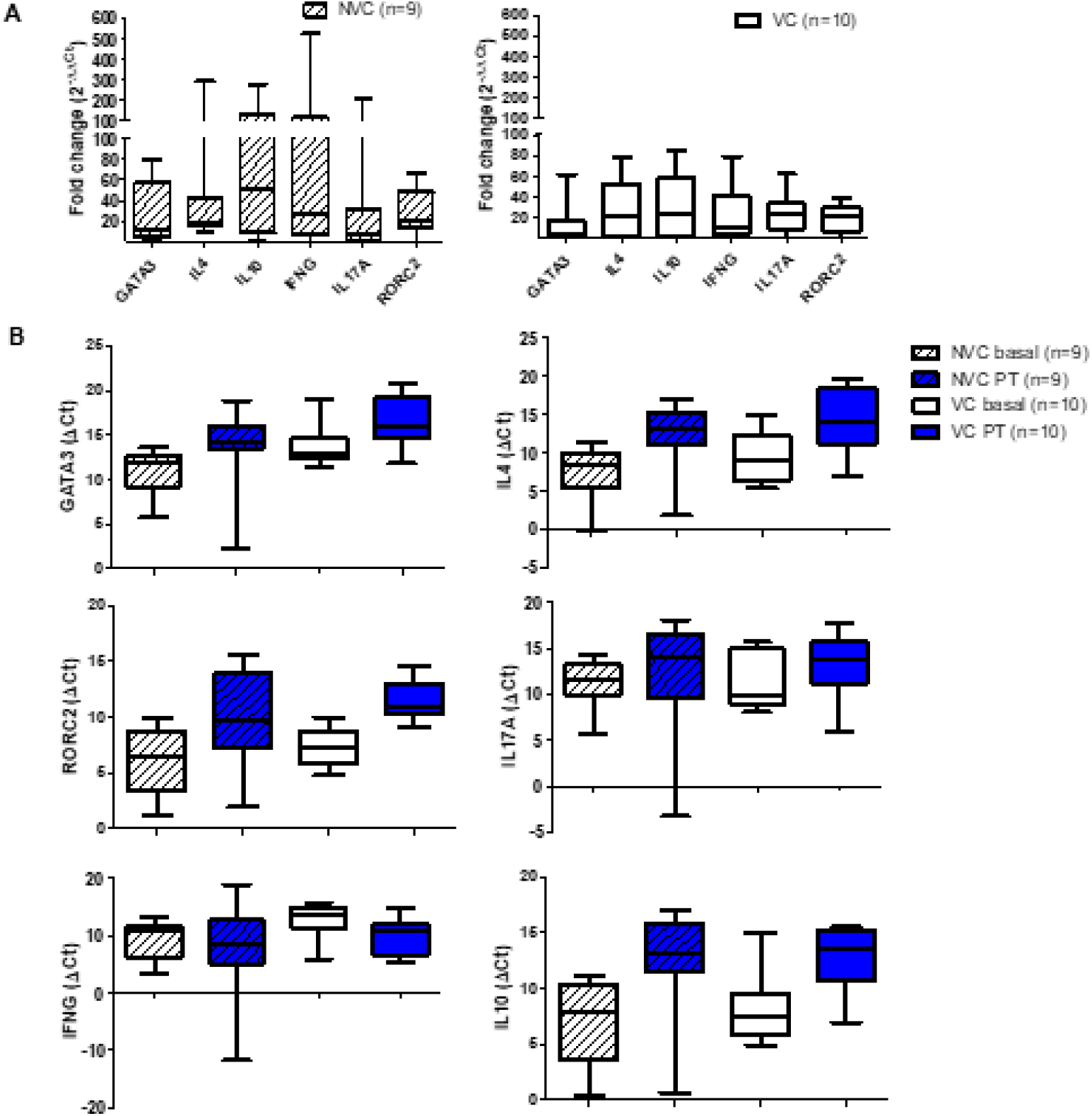
Cytokine expression in PT-stimulated PBMCs from infants born either from vaccinated (VC) or nonvaccinated (NVC) mothers. (A) Relative gene expression is similar in children born to Tdap-vaccinated (VC) or nonvaccinated (NVC) mothers when culturing PMBCs with PT for 120 h. Box represents variation between 25th and 75th percentiles, and whiskers represent minimum and maximum values. (B) VC presented higher basal expression of *GATA3* and *IFNG* than NVC, though differences disappear with PT stimulation for 120 h. Box represents variation between the 25th and 75th percentiles, and whiskers represent minimum and maximum values. Statistically significant differences are indicated.

In Table 2 it shown the total quantification from detectable all cytokines but IL-9 and IL-17F, as those values were undetectable.

Overall, TNF, IFN-γ, IL-13 and IL-6 were the most produced cytokines in both groups, with IL-6 being more produced in PT-stimulated cells derived from children born to vaccinated mothers (VC). In contrast, IL-2 was higher is children whose mothers were not vaccinated (NVC).

As both groups showed simultaneous IFN-γ and IL-13 production, which could indicate a mixed Th1/Th2 response or a good response by individuals to all cytokines, we evaluated the correlation between cytokines from different profiles, looking for a predominant response in all individuals or whether some individuals were indistinctly good responders to all T helper cell profiles (data not shown). However, no correlation was found between groups or cytokines. Individual responses were different from each other, making it challenging to find a correlation in a small group.

Regarding infants with higher cytokine levels, we analysed whether factors such as Tdap dose, gestational age, route of birth, infant’s sex, period of DTwP vaccination and exclusive breastfeeding could influence the response deviation (data not shown), but no significant correlations were found.

## Discussion

This study was the first to show the cellular immune response in older children, after the primary pertussis vaccination series, born to Tdap-vaccinated mothers in Brazil. Despite the use of Tdap during pregnancy was implemented in Brazil and other countries in the last few years and substantial data regarding antibody transfer and its influence on children’s humoral response to vaccination [10,26–28], little is known whether this strategy could affect the infant’s cellular immune response.

A few studies have been published recently about the influence of maternal antibodies in the neonatal humoral response, as the study made by Ibrahim et al. [29]. Women that did not receive Tdap during pregnancy but had detectable pertussis antibody levels showed that maternal antibody titres at delivery did not blunt infant vaccine response among children that receive at least 2 wP doses, and the authors observed that infant antibody titres increased with increasing maternal antibody levels against all pertussis antigens. A Brazilian study found that Tdap maternal immunization yielded significantly higher anti-PT IgG levels in vaccinated mothers and their infants compared to their unvaccinated counterparts and a strong positive correlation between the anti-PT IgG levels in maternal and cord blood at delivery, but did not look for antibodies against other pertussis antigens [10].

Lima et al. [28] showed the same correlation regarding antibodies against other bacterial antigens, and showed for the first time the analysis of the cellular immune response in whole cord blood cells, showing IFN-γ, IL-6, IL-10 and TNF production by neonatal cells upon stimulation with the whole inactivated *B. pertussis*, and low levels of IL-2, IL-4 and IL-17A.

In our study, we complemented the neonatal data by evaluating the cytokine productions in older children, with the complete primary series of vaccination against pertussis. The cytokines analysed were TNF-α and IL-2, which belong to a Th1 profile, in association with IFN-γ; IL-9 by Th9 cells, IL-4, IL-5 and IL-13, which contribute to the Th2 response profile; IL-10 is secreted by T regulatory cells and IL-17A, IL-17F, IL-21 and Il-22 secreted by Th17 cells. IL-6 is an inflammatory cytokine and is important in the differentiation of Th17 cells [24].

We used purified PBMC stimulated with PT, for five days, as described in the literature [30]. By using PBMCs it easier to access lymphocytes present in the samples and PT is the only specific antigen of *B. pertussis* [31], so our group decided to first run the assays with this antigen to have a first glimpse of what could have been happening, as well as, with PT, we analysed the adaptive response directly related to this stimulation. We could not determine the purity of PT used in the assays, but even with other bacterial antigens present we could see cytokine production upon stimulus.

Despite using samples from older children, our study agrees with Lima et al. [28] in the production of IFN-γ, IL-6 and TNF, which we found elevated in infants’ samples, but not IL-10. We also found low levels of IL-4 and IL-17A, but IL-13, which is an indicative of the Th2 response and was not available in the assay the authors performed, was found elevated in our samples. In all cytokines evaluated, there was one or more individuals that we did not detect any production, indicating a strong variability amongst human samples. As our both groups presented similar characteristics, the slight difference in gene expression could suggest modulation by vaccination. Even so, samples were limited, and the mRNA analysed was just to have a glimpse of what cells populations may have been present upon stimulus. Even though cytokine mRNA are produced earlier under these experimental conditions, the presence of the genes can show a tendency to a certain profile. This is seen in both the cytokine measurement and gene expression and could be complemented by the analysis of *TBX21*, the gene responsible for Th1 cell polarization [32].

When attempting to find correlation between different cytokines production, we found that in both groups every individual produced high levels of at least one cytokine. In the NVC group, a correlation between IFN-γ and IL-10 could point to a balance between activation and regulation, while in the VC group the same correlation was found, as well as between and IFN-γ and IL-13 (data not shown). This could indicate a mixed Th1/Th2 profile, with IL-10 possibly balancing the production of IFN-γ and the activation of Th1 cells. In both groups, cytokine levels were variable between individuals, and thus, it is not possible to determine a predominant response pattern.

According to Li *et al.* [33], Th2 cells do not prevent Th1 polarization but induce their apoptosis. In addition, they also observed that IL-10 could promote Th1 cell anergy through negative regulation of accessory molecules or IL-12 in antigen-presenting cells. Even inducing protection, maternal antibodies can also interfere with the child’s vaccine response, reducing the vaccine efficiency [34], leading to a decrease in the child’s antibody production via immune complex formation, antigen elimination via phagocytosis or epitope masking [35]. However, immune cellular response appears to be unaffected by these mechanisms [26]. As in Brazil children are vaccinated with DTwP, unlike many countries, we aimed to analyse the cellular immune response of vaccinated children born to either vaccinated or non vaccinated women, to confirm whether the cellular response would be intact. No difference was found between the two groups. Regardless of the different sample sizes and individual variabilities, both common factors in human vaccine studies [36–38], we could see IFN-γ and TNF-α production, indicative of the Th1 response, while only IL-13 from the Th2 profile was detected.

Overall, this work is the first step towards a complete scenario of maternal antibodies interference in cellular immune responses. Most pertussis studies describe humoral responses, before [39] and after the implementation of maternal vaccination [40,41], evaluating immunoglobulin levels produced by mothers and transferred to newborns, in countries using aP vaccines in children, unlike Brazil. A paper in Argentina describes the use of DTwP in children in the country; however, the study shows only the influence of maternal antibodies on the humoral responses of children before DTwP vaccination [41]. Despite the limitations, we evidenced cellular immune response in the context of pertussis vaccination, which is shown to be important for protection [42]. Our results show no difference in cytokine production against a bacterial antigen between groups, regardless of maternal vaccination, and this could set the path to other works that will support public health policies regarding vaccination in pregnancy. This knowledge is essential to the enhancement of vaccine protocols in pregnant women and as a basis for physicians and healthcare managers to recommend this strategy.

## Conclusion

This work shows cytokine production against a pertussis antigen in the context of maternal antibodies, and data could be strengthened by verifying these observations with more than one pertussis antigens. Also, there are ongoing studies that measured antibody responses to infant immunisation, in order to complement data and promote discussion about a potential difference in the impact of maternal immunisation of infant cellular and humoral responses.

## Acknowledgments

The authors would like to thank the teams at Hospital Maternidade Interlagos and Hospital Maternidade Leonor Mendes de Barros for recruitment and their help with blood specimen collection, Laboratory of Dermatology and Immunodeficiencies (LIM-56, Medical School of the University of São Paulo) and Immunology Centre from the Adolfo Lutz Institute for the infrastructure and support, the funding sources and all participants of the study. This study was supported by funding provided by the Special Health Fund for Mass Immunization and Disease Control (FESIMA) and São Paulo Research Foundation (FAPESP), grants #2015/19726-7 and #2015/16157-1. The funders had no role in study design, data collection and interpretation, or the decision to submit the work for publication. The authors also declare no conflicts of interest to disclose.

